# A cross-species meta-transcriptomic analysis of viral infection-induced shifts in microbial transcript profiles across insects

**DOI:** 10.64898/2026.09.28.754049

**Authors:** Adwait Mishra, Triveni Shelke, Vanika Gupta

## Abstract

Microbiome has co-evolved with insects over millions of years, establishing symbiotic relationships that influence their fitness and shape evolutionary trajectories. These long-term associations are shaped by the host’s ecology, diet, and physiology which determines microbial diversity. In turn, external stresses such as pathogenic infections can adversely affect the host microbiome, further compounding the effects of pathogenic infections on the host. Using a computational pipeline and publicly available transcriptomics datasets, we analyzed the effect of viral infections on transcript derived microbiome diversity and composition in various insects including *Drosophila melanogaster*, *Acyrthosiphon pisum*, *Antheraea pernyi*, and *Culex pipiens*. The meta-transcriptomic reanalysis provides an indirect but comprehensive comparison of infection-associated transcriptional microbiome changes across hosts with distinct life histories. Our study reveals that viral infections influence the host-associated microbial transcripts, leading to shifts in transcript-derived microbial profiles and offering insights into host-microbe dynamics. In *Cx. pipiens*, microbial transcript profiles remained relatively stable following RVFV exposure and were dominated by the bacterial endosymbiont *Wolbachia*. Our study highlights viral infections as key drivers of infection-associated shifts in microbial profiles and suggests a potential association between *Wolbachia* dominance and transcriptional microbiome stability in insects, with age as a contributing factor in shaping microbial transcript profiles in *Culex*. This cross-species framework uncovers conserved and host-specific patterns of virus-microbiome interactions, expanding our understanding of microbial resilience mechanisms in insects.

**Graphical abstract:** 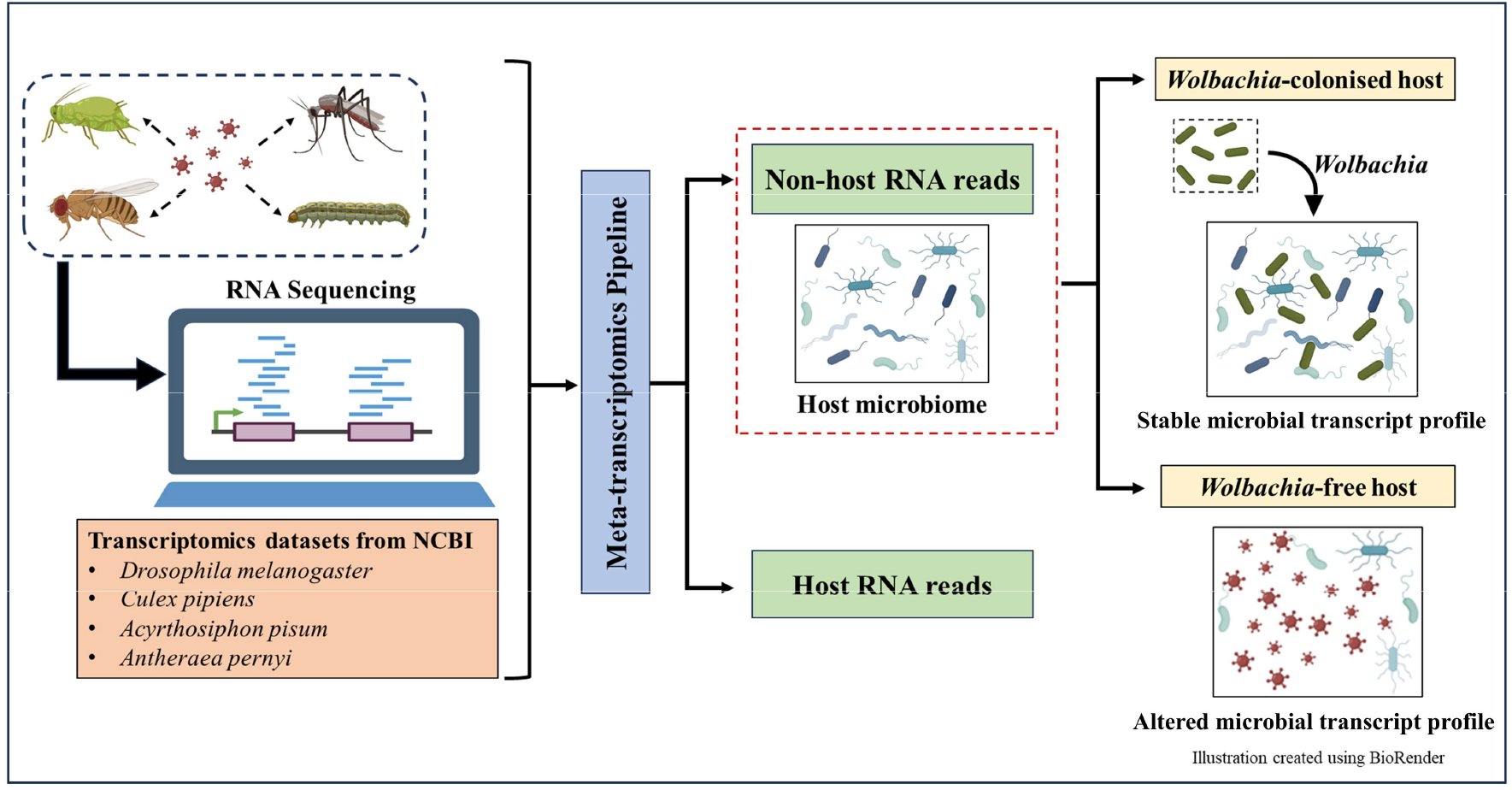

## 1. Introduction

Insects and their microbiomes have coexisted and evolved over millions of years, resulting in complex and often specialized symbiotic relationships. These interactions between insects and their microbiomes can be mutualistic, pathogenic, or neutral, with each influencing host fitness in distinct ways. Among these relationships, the most vital is symbiosis between insects and their gut microbiota, which plays a crucial role in improving immune function, digestion, and overall insect physiology [1–3]. Various insect species represent diverse ecological and physiological systems that shape their microbiome. The microbial composition is not static; it evolves over time due to various factors, including the host species [4], developmental stage [5], aging [6,7], host immunity [8–10], infections by other pathogens [11,12], and host metabolic status [13]. The mosquito *Culex pipiens* is a hematophagous insect whose microbiome composition changes drastically with blood feeding and oxidative stress [14]. The fruit fly *Drosophila melanogaster* harbors a simple gut community dominated by *Acetobacter* and *Lactobacillus*, influenced by diet and developmental stage [15,16]. The pea aphids *Acyrthosiphon pisum* maintains an obligate symbiosis with *Buchnera aphidicola*, essential for amino acid synthesis [17]. The silkworm *Antheraea pernyi* possesses a gut microbiome influenced by its exclusive oak leaf diet and larval development, with immune responses linked to microbial shifts during infection [18,19]. These ecological and physiological differences profoundly influence microbial resilience, emphasizing the importance of host context in interpreting cross-species microbiome patterns [20]. In addition to the direct benefits provided by these microbial communities, they also influence broader ecological and evolutionary processes, including population dynamics and species distribution [20].

The microbiome’s role in maintaining the host’s health becomes particularly significant when pathogenic infection weakens the host’s defenses [21]. Pathogenic infections can disrupt gut homeostasis inducing microbial dysbiosis - an imbalance in the gut microbiome, that in turn can adversely affect host health. Further, microbial dysbiosis can compromise the immune system, leaving hosts more susceptible to infections [22,23]. Infection of mosquitoes with *Beauveria bassiana* induced microbiome imbalance, highlighting the role of pathogens in shaping the insect microbiome [24,25]. In addition of extracellular pathogens, intracellular endosymbionts such as *Wolbachia* have also been associated with alterations in gut microbial community structure. In *D. melanogaster*, *Wolbachia* correlates with reduced abundance of *Acetobacter*, despite *Wolbachia* being absent from the gut lumen and therefore unlikely to interact directly with resident gut bacteria [26]. The absence of detectable changes in AMP’s expression or other canonical immune effector responses implies that *Wolbachia*-associated microbiome changes arise indirectly as a consequence of complex host physiological perturbations such as gut pH, rather than direct immune regulation or microbe-microbe interactions [26,27].

In the context of viral infections, intestinal bacteria enhance the replication and transmission of several enteric viruses (e.g., poliovirus, murine norovirus, reovirus, rotavirus) through direct interactions between bacterial surface molecules such as lipopolysaccharide and viral particles, which stabilize virions and promote viral attachment to host cells [28]. The virome, the viral component of the microbiome, can influence the host by stimulating basal antiviral immune signaling [29] and maintaining intestinal homeostasis, as seen with murine norovirus, which mimics commensal bacteria [30]. However, these virus-microbiota interactions are highly contingent on microbial species composition and diversity [31]. A deeper understanding of these virus-microbiome dynamics in insect hosts is therefore essential for advancing insect health, pest management, and strategies to mitigate virus transmission.

The specific effects of viral infections on insect microbiomes remain poorly understood, and this study addresses that gap using a cross-species meta-transcriptomic approach. While numerous transcriptomic studies have investigated host gene expression responses to viral infection, they often overlook microbial community composition and fail to capture infection-associated transcriptional microbial dynamics that may be present within these datasets [32–35]. Notably, the unmapped reads from such RNA-seq datasets can contain valuable information about the microbiome and the virome. Several studies have demonstrated that transcriptomic datasets can be used to characterize host-associated microbial communities, and investigate functional microbial dynamics [36–38], highlighting the utility of RNA-seq data as a valuable resource for microbiome research beyond host gene expression analyses. To capture microbiome responses across diverse ecological contexts, we selected four phylogenetically distinct insect hosts representing different feeding habits: *D. melanogaster* (frugivorous) [32], *Cx. pipiens* (hematophagous) [33], *A. pisum* (phytophagous) [35], and *A. pernyi* (folivorous) [34]. Feeding behavior is known to strongly influence insect-associated microbial communities; therefore, selecting hosts with different diets allows broader comparison of microbial response to viral infection. In addition, each of these insect species was infected with a different virus, enabling cross-species examination of virus-associated microbial responses. The datasets were further selected based on the availability of matched control and virus-infected samples with at least three biological replicates per condition, sufficient sequencing depth, and inclusion of wild-type insects to avoid genetic biases. Although most datasets represented whole-body RNA-seq samples, the A. pernyi dataset comprised dissected larval cuticle tissue generated to investigate cuticle liquefaction following viral infection. Together, these criteria ensured both comparability across conditions and broad biological relevance across host systems. In this study, we implemented a computational pipeline that reanalyzes these unmapped reads, aligning them to non-host microbial and viral references to reconstruct transcript-derived microbial and viral profiles from the same sequencing run. This approach enabled us to investigate how viral infections reshape microbial transcripts representation and how host-symbiont context modulates these effects. Although RNA-seq is not specifically designed for microbiome profiling, our meta-transcriptomic framework effectively captures microbial transcript signals from host samples and reveals ecologically meaningful patterns-such as infection-associated shifts in transcript derived microbial profiles, taxon-specific responses, and symbiont-mediated effects-providing a cost-effective and integrative view of host-virus-microbe interactions. Overall, our results shed light on the dynamics of virus-induced shifts in transcriptional microbial patterns across diverse insect hosts and emphasize the untapped potential of transcriptomic data to reveal microbiome dynamics.

## 2. Methodology

### 2.1. Data extraction and quality assessment

Dataset selection was guided by sequencing quality and biological diversity, ensuring inclusion of insects with diverse feeding habits, life histories, and symbiotic associations to capture a broad spectrum of virus–microbiome interactions. RNA-seq datasets were retrieved from the National Center for Biotechnology Information (NCBI) database and only Illumina-based transcriptomic datasets generated following viral infection were included (Table 1). Importantly, these datasets encompass insect species representing diverse ecological and physiological systems, including frugivorous, hematophagous, phytophagous, and folivorous hosts. Moreover, these datasets represented both systemic and oral viral infection models, enabling a broader assessment of infection-associated transcriptional microbial dynamics. This diversity facilitated assessment allowed us to evaluate the robustness and generalizability of our meta-transcriptomic framework across different host–virus contexts and environmental niches.

**Table 1.** List of RNA-seq datasets used for meta-transcriptomic analysis.

| Dataset | Bioproject-ID | Host | Infection | Feeding habits | Source of RNA | NGS | References |
| --- | --- | --- | --- | --- | --- | --- | --- |
| **Dm** | PRJNA682591 | *Drosophila melanogaster* | Sindbis virus (SINV) | Frugivorous | Adult whole body | NextSeq 500 | [32] |
| **Cp** | PRJNA596664 | *Culex pipiens* | Rift valley fever phlebovirus (RVFV) | Hematophagous | Adult whole body | Illumina HiSeq 2000 | [33] |
| **Api** | PRJNA419454 | *Acyrtosiphon pisum* | Acyrtosiphon pisum virus (APV) | Phytophagous | Adult whole body | Illumina HiSeq 4000 | [35] |
| **Ape** | PRJNA517653 | *Antheraea pernyi* | Antheraea pernyi nucleopolyhedrovirus (ApNPV) | Folivorous | 5<sup>th</sup> instar larval cuticle tissue | Illumina HiSeq 4000 | [34] |

The raw reads were first subjected to quality assessment using FastQC (v0.11.9) [39], to obtain metrics related to per-base sequence quality. In particular, the per-base quality score profiles (Phred scores) were examined to ensure that the majority of bases exceeded the Q30 threshold, indicating high sequencing accuracy. FastQC also flagged the presence of adapter contamination. When adapter content or low-quality bases were detected, reads were processed using Fastp (v0.23.2) [40], for adapter trimming and quality filtering to ensure clean, high-quality sequences for downstream analyses.

### 2.2. Sequence alignment and processing

These high quality reads were aligned to the reference genome for each host using STAR (v2.7.10b), which works by finding the longest matching sequence (Maximal Mappable Prefix, MMP) between each read (or read pair) and the reference genome, and detects spliced junctions as well as fusion reads [41]. Reference genome assemblies were obtained from the NCBI, following genome assemblies were used for alignment: *D. melanogaster* (RefSeq accession GCF_000001215.4), *A. pernyi* (GenBank accession GCA_015888305.1), *A. pisum* (RefSeq accession GCF_005508785.2), and *C. pipiens* (RefSeq accession GCF_016801865.2). Following alignment, the coordinate-sorted BAM file (Aligned.sortedByCoord.out.bam) generated by STAR were used as the starting point for the meta-transcriptomics analysis pipeline. This BAM file contain detailed alignment information, including mapping coordinates, alignment flags, and read quality metrics, allowing discrimination between reads that aligned to the host genome and those that remained unmapped. We implemented a meta-transcriptomics pipeline (Fig. 1) designed to distinguish between host and non-host RNA reads [42]. This framework allowed us to extract transcriptional microbial and viral signals from standard host RNA-seq data, enabling integrative analysis of host-microbiome interactions. The subsequent computational steps were performed:

**Fig. 1.**
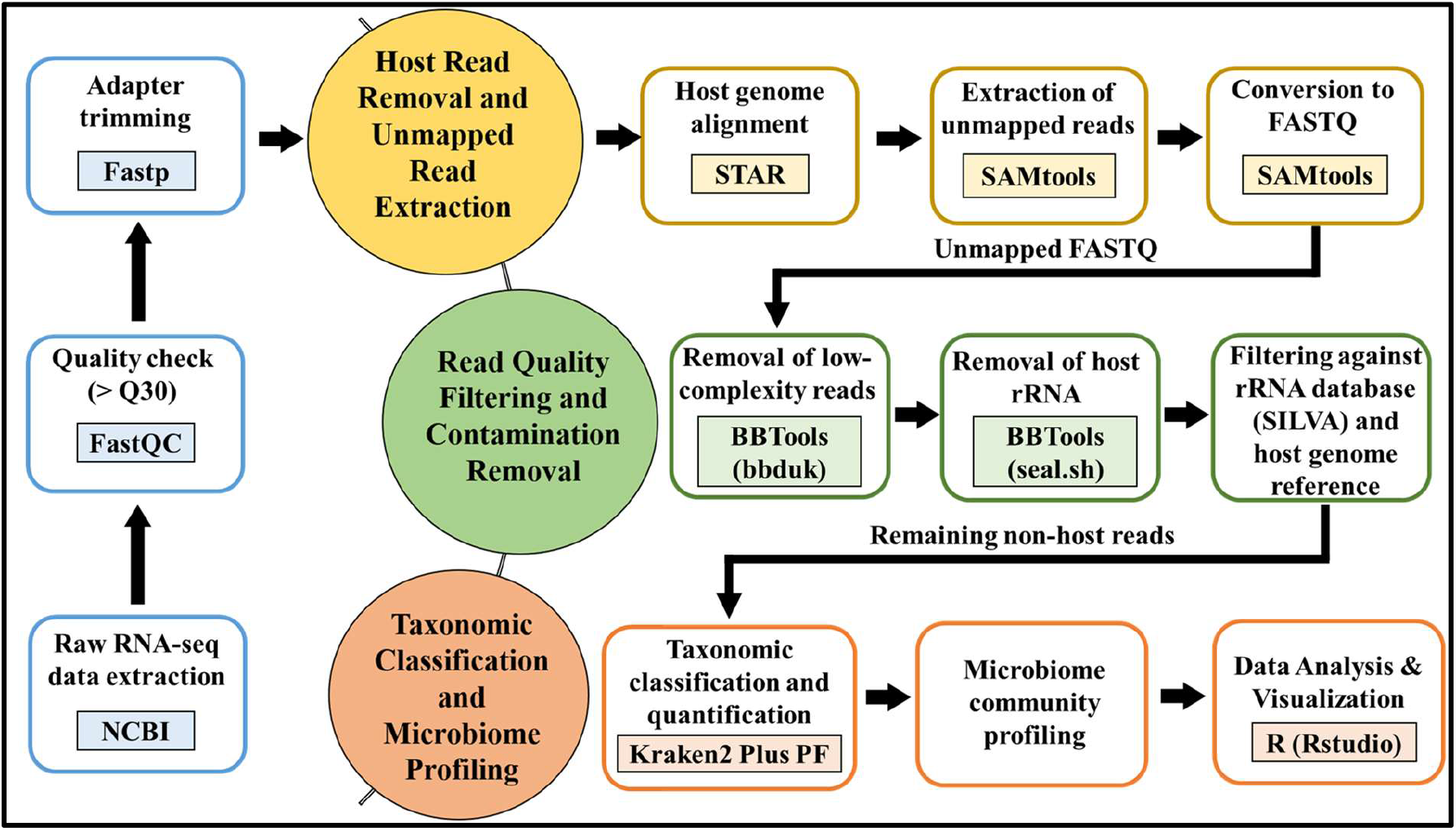
Workflow of the meta-transcriptomic pipeline including quality control, host alignment, extraction of unmapped reads, rRNA filtering, taxonomic classification.

#### 2.2.1. Extraction of Unmapped Reads

To obtain non-host reads, unmapped sequences were first isolated from the coordinate-sorted BAM files using SAMtools (v1.16) by selecting reads that failed to align to the host genome (-f 4 flag) [43]. Unmapped reads in the datasets may originate from non-host sources such as bacteria, viruses or fungi, and thus would be useful for studying microbial or viral sequences present in the sample. The resulting BAM file containing unmapped reads was subsequently converted to FASTQ format for downstream analyses. These unmapped reads formed the basis for subsequent non-host taxonomic profiling.

#### 2.2.2. Quality control and trimming

The unmapped reads were further processed to remove low-complexity sequences using tools from the BBTools (v37.62) [44]. Specifically, low-complexity reads dominated by simple or repetitive sequences (e.g., poly-A or homopolymeric regions) were filtered using bbduk.sh (with entropy=0.5, entropywindow=50 and entropyk=5). As a result, only high-complexity reads were retained for downstream taxonomic classification.

#### 2.2.3. Ribosomal RNA (rRNA) and host contamination Filtering

To enhance taxonomic classification specificity, rRNA and residual host-derived sequences were removed using the seal.sh script from the BBTools suite. rRNA molecules are highly abundant in RNA sequencing libraries and may remain present even after experimental rRNA depletion, potentially dominating sequencing reads and reducing the detection of informative microbial transcripts. First, unmapped reads were screened against host rRNA references to remove insect-derived rRNA sequences. Reads matching the reference were separated, while non-matching reads were retained for further analysis. The remaining reads were screened against the SILVA rRNA database (release 138; LSURef and SSURef datasets) as well as the host genome reference. This step removed residual rRNA sequences from diverse organisms as well as any remaining host genomic contamination. Reads that did not match either reference was retained as high-confidence non-host sequences for downstream microbial taxonomic classification.

#### 2.2.4. Taxonomic classification

The Cleaned and filtered reads were classified using Kraken2 (v2.1.2). Kraken2 compares k-mers from sequencing reads to a pre-built database that includes bacteria, archaea, viruses, and the fungal genome [45]. In this study, classification was performed using the Kraken2 PlusPF database, which provides broad coverage of microbial genomes. Reads were classified using a confidence threshold of 0.3, Kraken2 generated both detailed classification outputs and summary reports for each sample, including the number and percentage of reads assigned to different taxonomic groups as well as unclassified reads. These reports were subsequently used to characterize the microbial and viral composition at transcriptome level associated with each host sample and to investigate potential infection-associated transcriptional microbial shifts. Taxa were reported at the lowest taxonomic level assigned by Kraken2. Consequently, some microbial taxa were resolved to species level, whereas others were classified only to the genus level due to limitations in sequence resolution or database matches. As the microbiome was inferred from unmapped reads of host RNA-seq datasets, the resulting profiles represent transcriptionally active microbial sequences rather than absolute microbial abundance. Therefore, diversity and abundance metrics in this study reflect transcript-derived representation of microbial taxa rather than direct measurements of microbial population size.

### 2.3. Microbiome analysis

#### 2.3.1 Relative transcript abundance and diversity estimation

Raw read counts obtained from Kraken2 classification were normalized to relative abundance to account for differences in sequencing depth among samples. This normalization reflects proportional representation within the detected transcript pool and does not account for absolute microbial load. Specifically, taxon counts within each sample were converted to percentages of the total classified reads per sample using the transform_sample_counts function in the ’phyloseq’ package in R (version 4.3.1). Relative abundance was calculated by converting taxon read counts into percentages of total classified reads per sample. Taxa with relative abundance >1% in at least one sample were retained for downstream analyses. Transcriptional alpha diversity across samples was quantified using the Shannon index [46], calculated with the ’phyloseq’ package.

#### 2.3.2 Statistical analyses

To assess differences in microbial diversity at transcriptome level between control and infected samples in all four datasets, we performed unpaired t-tests. For *Culex*, we also examined the effect of age and infection status transcriptional microbiome composition using a two-way ANOVA. The following model was used:

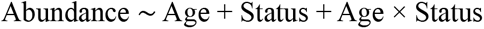

Here, abundance (relative transcript abundance of a given microbial taxa) was the response variable, while Age and Infection status were categorical predictors, with their interaction term included to test whether the effect of infection varied across age groups. This model was applied separately to test-*Wolbachia* abundance (Table 4), overall transcriptional microbiome composition (Table 5), and other microbial taxa excluding *Wolbachia* (Table 6).

Principal Component Analysis (PCA) of microbial transcriptome was conducted to reduce dimensionality and summarize variance across samples to evaluate overall community composition. The first two principal components (PC1 and PC2) were extracted for visualization. Because microbial communities varied significantly among host species, the combined PCA was only used to show overall patterns at the host level. It was not used to find similarities related to infection across species. To statistically assess differences in community composition (transcriptional beta-diversity) between control and infected samples, we applied Permutational Multivariate Analysis of Variance (PERMANOVA) using the vegan package [47], based on Bray-Curtis dissimilarity matrices calculated from normalized transcript abundance data. GLMs were implemented to test associations between explanatory variables (infection status and host species) and microbiome alpha diversity (Shannon index).

#### 2.3.3 Data visualization

All boxplots and PCA plots were generated using ’ggplot2’ [48]. Heatmaps were created with ’pheatmap’ [49]. For heatmap visualization, relative transcript abundance values were log10-transformed after adding a small pseudocount (0.01) to stabilize variance and improve interpretability of low-abundance taxa.

## 3. Results

### 3.1 Viral infection alters transcriptional microbiome diversity in insects

For our study, we have utilized four publicly available RNA-seq datasets of distinct insect hosts. These include Dm (*D. melanogaster*) infected with SINV, Cp (*Cx. pipiens)* exposed to RVFV, Api (*A. pisum*) infected with APV, and Ape (*A. pernyi*) infected with ApNPV. Notably, *A. pernyi* dataset was generated from dissected larval cuticle tissue rather than whole-body samples and therefore represents tissue-associated microbial transcripts specific to the cuticle environment (Table 1). Together, these datasets include phylogenetically and ecologically diverse hosts, providing an opportunity to compare infection-associated changes in transcript-derived microbial profiles across insects.

We first analyzed the microbial transcriptional Shannon diversity of our datasets to understand whether viral infection leads to shifts in transcript-derived microbial profiles. Hereafter, microbiome composition and perturbations refer to relative transcript-derived profiles of detected taxa. We found that datasets Dm, Api, and Ape exhibited a significant decrease in Shannon diversity index (p < 0.0001; Fig. 2, 3), suggesting an infection-associated reduction in the richness of detected microbial transcripts. In Dm, this effect was consistent across all three time points examined (6 hr, 24 hr, and 48 hr), each of which showed a significant loss of microbial transcriptional diversity relative to controls (Fig. 2a). In contrast, Cp showed no significant change in Shannon diversity at any time point: 2 hr (corresponding to 0 days), 3 days, or 14 days post-exposure (Fig. 2b). Notably, viral sequences were not detected above the classification threshold in this dataset using our pipeline (Supplementary Table S1). Both Rift Valley fever virus (TaxID: 11588) and Rift Valley fever phlebovirus (TaxID: 1933187) are included in the Kraken2 PlusPF database used in this study. This means that the lack of classified RVFV reads is not due to missing reference sequences in the database. This likely reflects a combination of poly(A)-based library preparation, which limits capture of non-polyadenylated viral transcripts such as RVFV, and the reduced sensitivity of k-mer-based classification for low-abundance viral sequences. Importantly, the original study corresponding to this dataset reported increased viral load in infected samples at all time points using RT-qPCR targeting RVFV-specific genes. Therefore, the absence of classified viral reads in our analysis should not be interpreted as absence of viral activity.

**Fig. 2.**
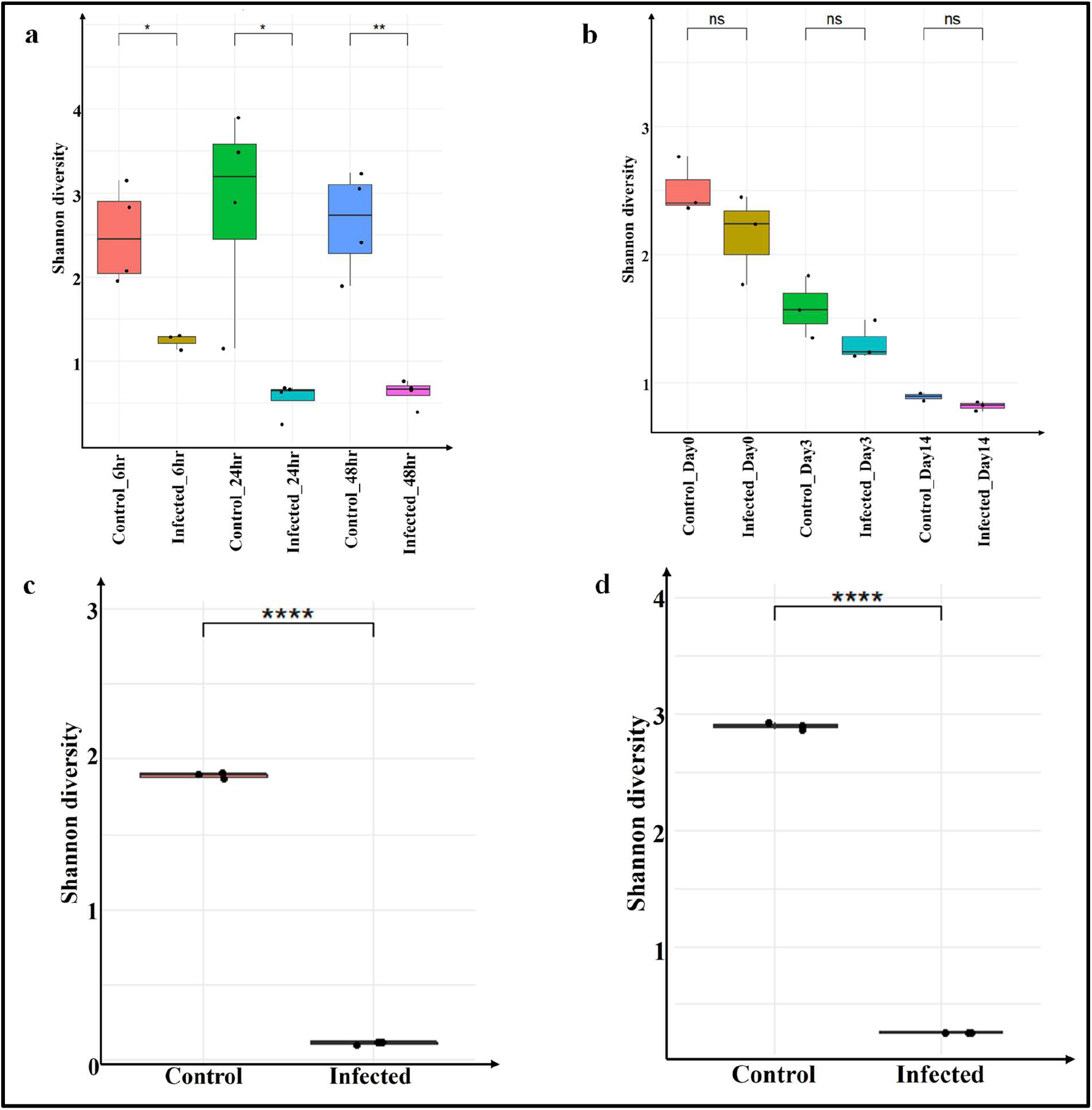
Estimation of transcriptional Shannon diversity index of microbiome richness across control and virus-infected samples in four insect species. **a** *Drosophila* (*Wolbachia* free) infected with SINV, showed significant declines in transcriptional microbial diversity at 6 hr, 24 hr and 48 hr post-infection. **b** *Cx. pipiens* infected with RVFV, showing no statistically significant differences in microbial transcripts across age-matched control and infected samples. **c** *A. pisum* infected with APV with a significant decrease in microbial transcripts in infected aphids. d *A. pernyi* infected with ApNPV also exhibited a significant reduction in microbial transcripts compared to controls. Statistical significance was assessed using t-tests; ns: non-significant; *p < 0.05, **p < 0.01, ***p < 0.001, ****p < 0.0001.

**Fig. 3.**
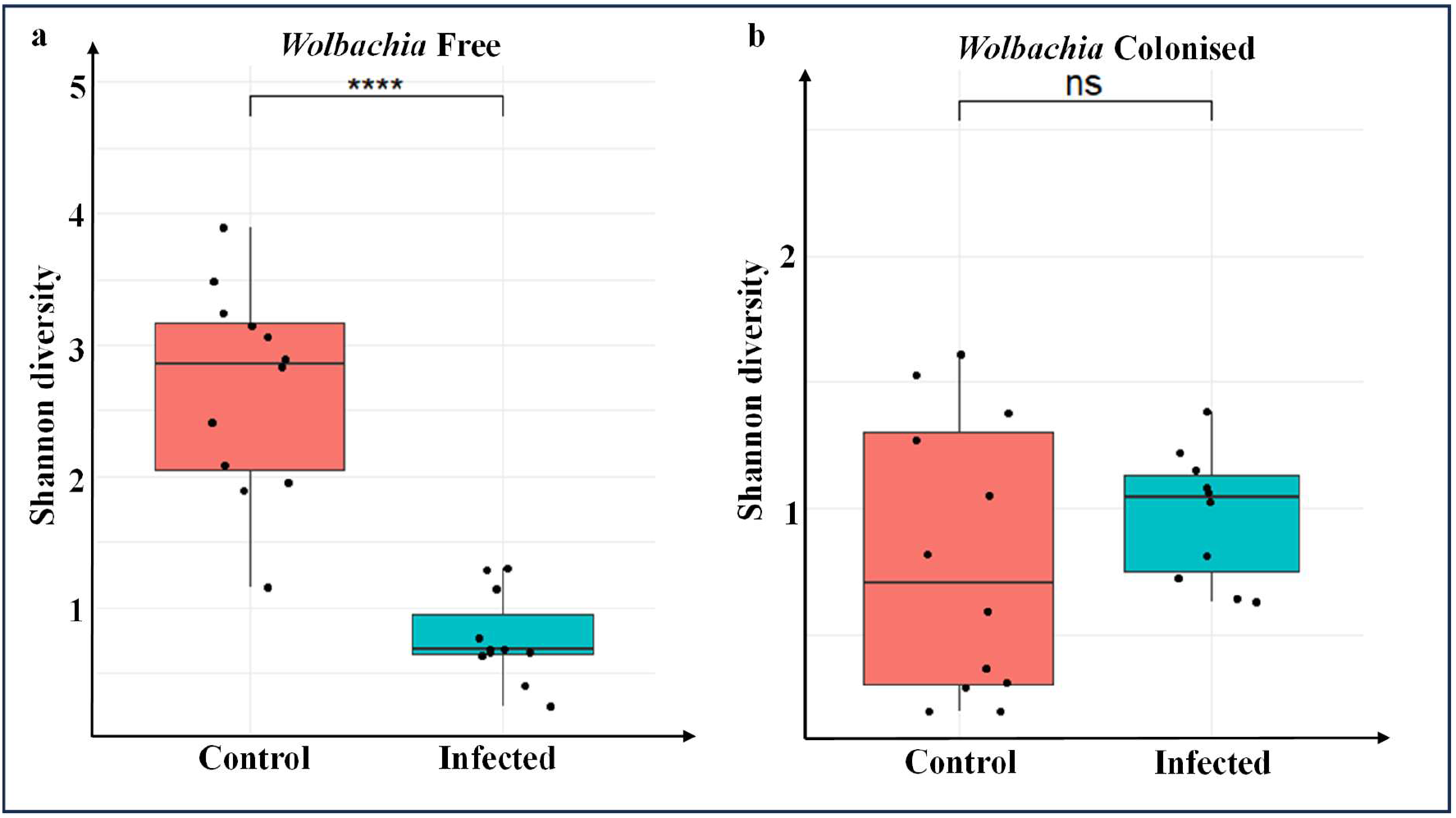
Boxplots representing transcriptional Shannon diversity index of the microbiome under viral infection. **a** *D. melanogaster* (*Wolbachia*-free) vs. **b** *D. melanogaster* (*Wolbachia*-colonized) infected with SINV. A significant reduction in microbial transcripts is observed in infected *Wolbachia-*free flies, whereas no significant difference is detected in *Wolbachia*-colonized flies, indicating a potential protective role of *Wolbachia* against infection-induced shifts in microbial transcriptome.

To examine overall transcriptional microbial community patterns across insects, we have generated heatmaps based on log-transformed relative transcript abundance (Fig. 4). This approach enables detailed, sample-wise comparison of transcriptional microbial profiles, capturing both dominant and low-abundance taxa. Three datasets (Dm, Api and Ape) show clear infection induced shifts in microbial composition, depicted by increase in viral transcript abundance following infection (Fig. 4a, 4c, 4d). In contrast, Cp shows no clear distinction in microbial transcript abundance between control and infected samples. However, it reflects age associated variation, with microbial transcript profiles remaining relatively stable and dominated by *Wolbachia* (Fig. 4b). We further examined shifts in transcriptional microbial community composition upon infection by comparing taxa with transcript relative abundances greater than 1% in infected vs. control samples. In Dm, the number of taxa with >1% transcript abundance declined from 40 in controls to 9 post-infection; whereas, in Api, number of taxa decreased from 6 to 2, with *B. aphidicola* being the only taxon exceeding 1% relative transcript abundance after infection. This pattern aligns with *Buchnera*’s obligate nutritional role, which likely ensures its persistence even during viral infection. Similarly, in dataset Ape, the number of taxa dropped from 12 in controls to 2 following infection. Interestingly, while the infecting virus accounted for the majority of relative abundance, iflavirus was the only virus among the other microbiome that consistently persisted in both control and infected samples in this dataset (Fig. 4d). The infecting virus occupying the majority of reads assigned to microbial taxa likely reflects both biological proliferation and transcript dominance.

**Fig. 4.**
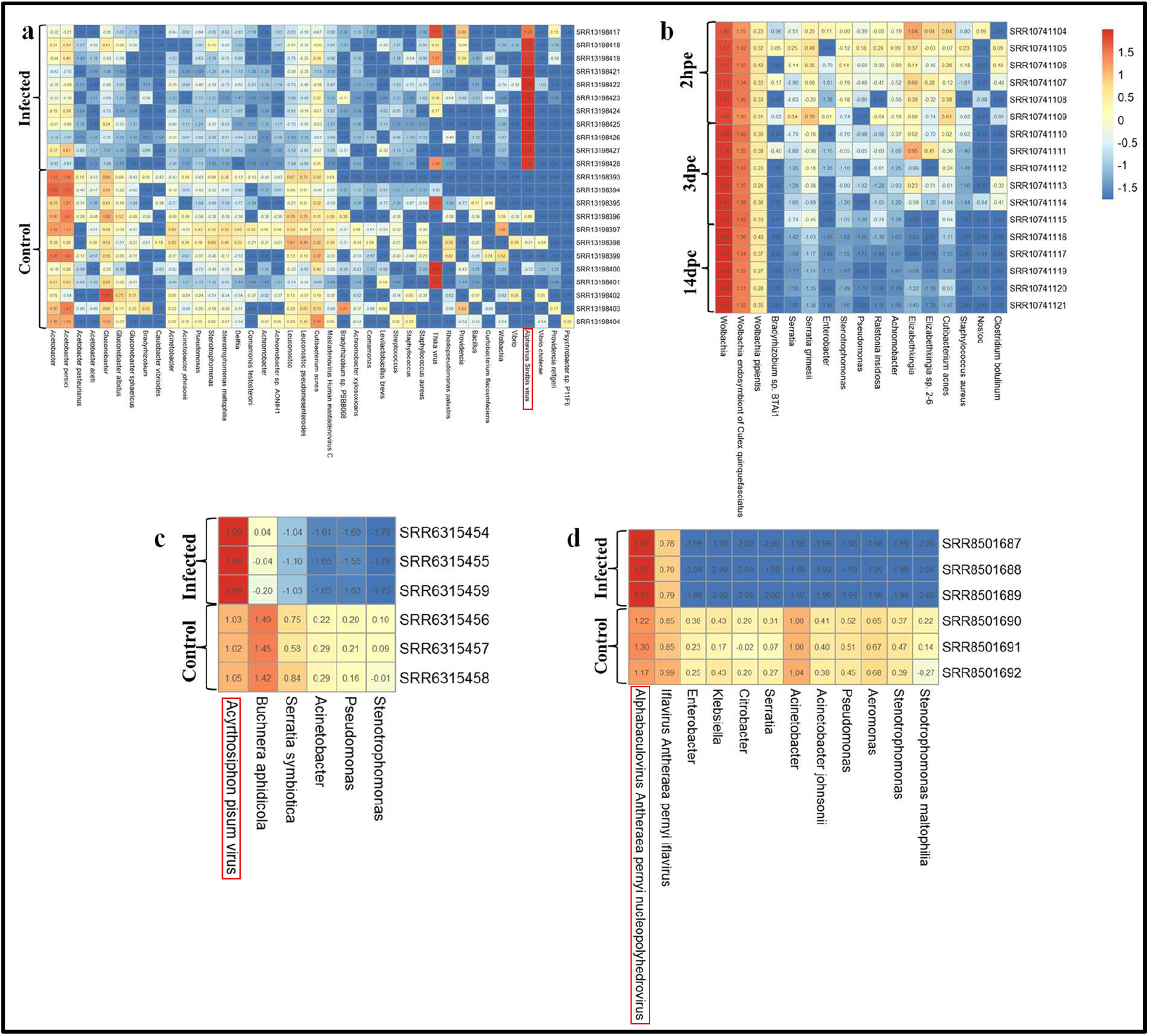
Heatmap of microbial community composition at transcriptome level based on log10-transformed relative transcript abundance (%), with taxa filtered to include only those exceeding 1% relative transcript abundance in at least one sample. **a** *Drosophila* (*Wolbachia* free) infected with SINV showing marked shifts in microbial transcript abundance post-infection across time points. **b** *Cx. pipiens* infected with RVFV highlighting age-associated variation with stable transcriptional microbial profiles dominated by *Wolbachia* post-infection (hpe-hours post-exposure, dpe-days post-exposure). **c** *A. pisum* infected with APV, where *B. aphidicola* remains the dominant taxa, despite infection-induced diversity loss. **d** *A. pernyi* infected with ApNPV, revealing a reduction in transcript-derived bacterial taxa with persistence of iflavirus in infected individuals. The color scale (approximately -1.5 to 1.5) reflects standardized log-transformed relative transcript abundances, rather than raw abundance values, with rows denoting individual samples and columns representing microbial taxa.

### 3.2 Comparative multivariate analyses reveal distinct clustering patterns depending on the infection status

PCA of microbial transcriptome was performed to examine the variation in transcriptional microbiome composition and to assess the impact of infection status (Fig. 5). Infection-associated clustering of microbial transcripts is evaluated within individual datasets, thereby avoiding overinterpretation of cross-species comparisons. The PCA plot revealed distinct clustering patterns for each species, suggesting potential host-specific and infection-associated effects on microbial community structure. For dataset Api and Ape, infected and control samples were separated along PC2, indicating potential differences in microbial transcript profiles between healthy and infected individuals for these hosts. In Dm, control and infected groups were distinguishable but positioned relatively close together on the PCA plot, suggesting a more moderate shift in microbial community composition associated with infection. For Cp, samples were tightly clustered with minimal spread and showed no apparent separation by infection status. This suggests a stable transcriptional microbiome composition that is not substantially altered by infection (Fig. 5).

**Fig. 5.**
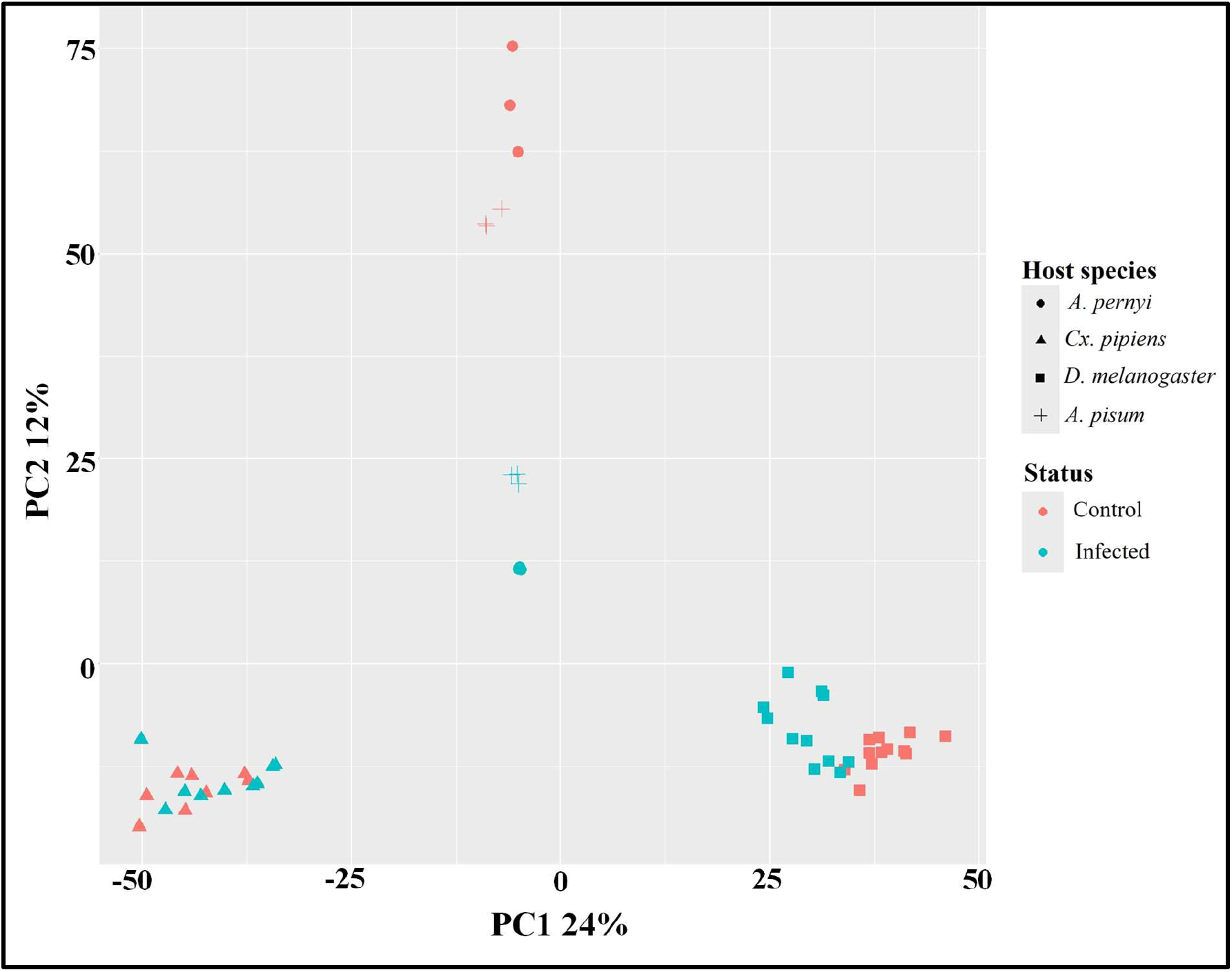
PCA ordination of microbial transcriptome showing variation in microbial community composition across host species and infection status. Each point represents the transcriptional microbial community structure of an individual sample. Distinct shapes indicate different host species. Different colors represent status. The first two principal components explain 24% (PC1) and 12% (PC2) of the total variance, respectively.

We conducted a PERMANOVA to test for differences in transcriptional microbial communities between control and infected samples (Table 2). Infection status was significantly associated with compositional changes in Dm (p = 0.001, R² = 0.12218), indicating a clear impact of infection despite moderate spatial separation in PCA. For Ape (p = 0.100, R² = 0.86067) and Api (p = 0.100, R² = 0.70809), no statistically significant differences were observed, despite high R² values. These higher p-values are likely due to limited sample sizes and the resulting constraints on the statistical power of permutation tests. In Cp, both PCA and PERMANOVA confirmed minimal compositional changes in microbial transcriptome due to infection (p = 0.249, R² = 0.07108). Together, these results indicate that the association between infection and transcriptional microbial composition may vary across host species.

**Table 2.** Results of the PERMANOVA assessing the effect of infection status on microbial composition at transcriptome level. ***p < 0.001.

| Host Species | Infection | R<sup>2</sup><br>(Model) | R<sup>2</sup><br>(Residual) | F<br>value | p-value |
| --- | --- | --- | --- | --- | --- |
| *D. melanogaster* | SINV | 0.12218 | 0.87782 | 2.92 | 0.001 |
| *Cx. pipiens* | RVFV | 0.07108 | 0.92892 | 1.14 | 0.249 |
| *A. pisum* | APV | 0.70809 | 0.29191 | 9.70 | 0.100 |
| *A. pernyi* | ApNPV | 0.86067 | 0.13933 | 24.70 | 0.100 |

### 3.3 GLM analysis supports infection-associated microbial transcripts changes

We employed a GLM to complement PERMANOVA and enhance the robustness of statistical evaluation, given the limited sample sizes. Unlike PERMANOVA, which assesses differences in overall community composition based on distance matrices, the GLM evaluates associations between explanatory variables and microbiome alpha diversity (Shannon index) across host species. The GLM analysis revealed a statistically significant association between infection and reduced transcriptional Shannon diversity in Ape (Estimate = −2.766 ± 0.0040, p < 0.001), Api (−1.840 ± 0.0290, p < 0.001), and Dm (−1.741 ± 0.2624, p < 0.001) microbiome (Table 3), suggesting that infection may be linked to decreased microbial transcript diversity. Conversely, infection did not significantly alter transcriptional microbiome composition in Cp (−0.304 ± 0.2904, p = 0.313), suggesting host-specific differences in microbiome response. These complementary analyses highlight that while PERMANOVA captures community-level compositional shifts based on transcriptional beta-diversity distances, the linear modelling framework allows testing whether infection status is associated with changes in transcriptional microbial diversity across datasets.

**Table 3.** GLM results for the effect of infection on insect species transcriptional microbiome. ***p < 0.001.

| Species | Estimate<br>± SE | t-<br>value | p-value | AIC | Residual<br>df | Residual<br>Deviance |
| --- | --- | --- | --- | --- | --- | --- |
| *D. melanogaster* | -1.741 ±<br>0.2624 | -6.634 | <0.001 | 47.82 | 21 | 8.30 |
| *Cx. pipiens* | -0.304 ±<br>0.2904 | -1.045 | 0.313 | 34.61 | 15 | 5.36 |
| *A. pisum* | -1.840 ±<br>0.0290 | -63.50 | <0.001 | -19.47 | 4 | 0.0050 |
| *A. pernyi* | -2.766 ±<br>0.0040 | -688.6 | <0.001 | -43.18 | 4 | 9.68×10<sup>-5</sup> |

### 3.4 *Wolbachia-*colonized insects maintain microbial transcript stability during viral infection

In both Dm and Cp, *Wolbachia*-positive insects showed no significant change in microbiome transcriptional Shannon diversity following viral infection (Fig. 2b, 3b). To further investigate the role of *Wolbachia* in shaping the *Culex* microbiome, we tested whether *Wolbachia* titre varies with mosquito age or infection status. Our two-way ANOVA on *Wolbachia* transcript abundance (Table 4) showed that *Wolbachia* transcripts increased significantly with age (p < 0.0001, Fig. 6), but viral infection had no effect. Further, no significant Age × Infection Status interaction was detected, highlighting that the age-dependent increase in *Wolbachia* transcripts is solely due to increasing age (Fig. 6). These age-dependent titer changes align with prior studies on *Wolbachia* dynamics during insect development [50,51].

**Fig. 6.**
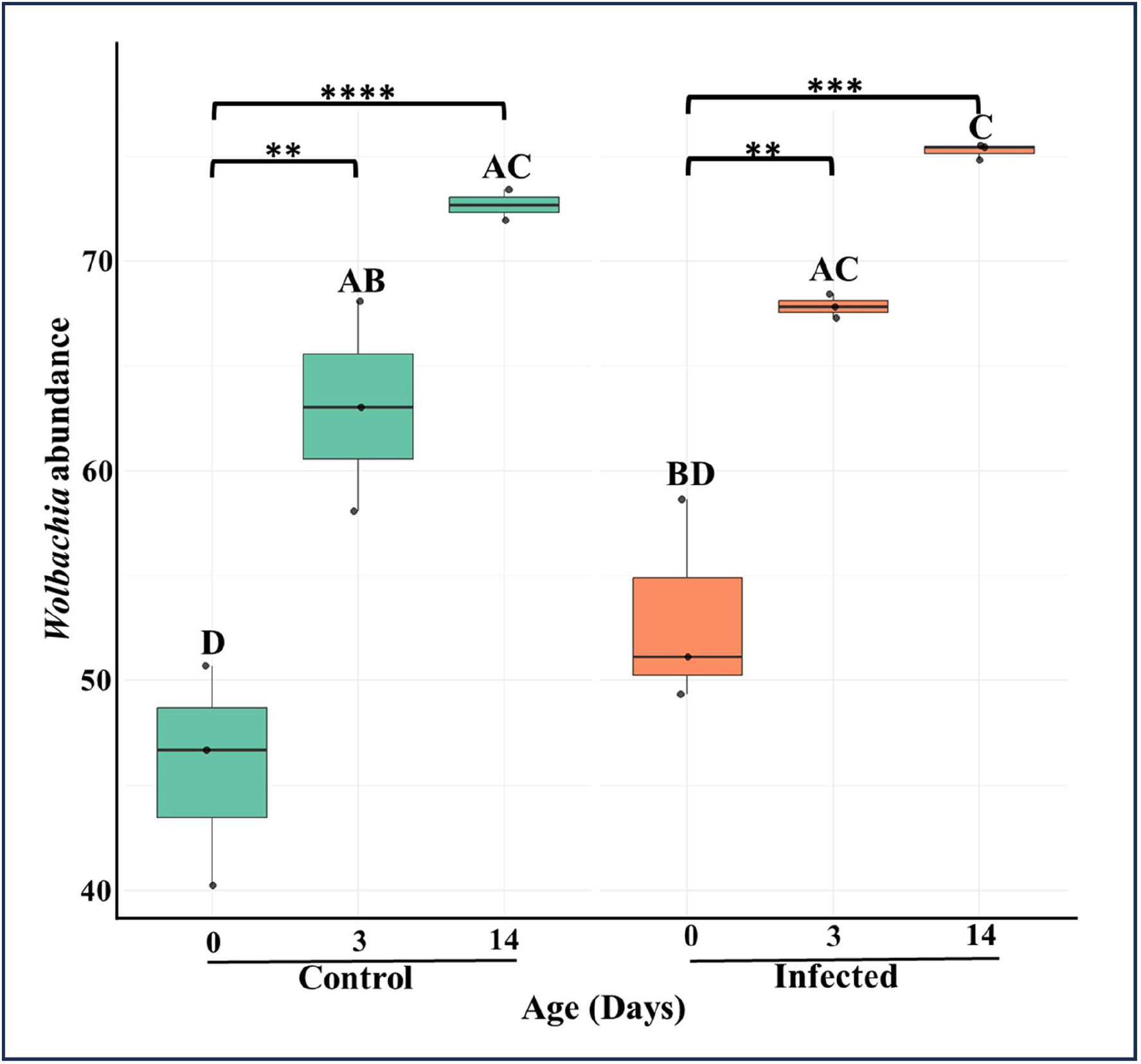
Boxplots showing *Wolbachia* transcript abundance in control and virus-infected *Cx. pipiens* at 0 days (corresponding to the 2 hr post-infection timepoint), 3 days, and 14 days post-infection. Letter annotations denote statistically distinct groupings based on Tukey’s post-hoc test, where groups sharing letters are not significantly different. Significance levels: *p < 0.05, **p < 0.01, ***p < 0.001, ****p < 0.0001. Each point represents an individual sample.

**Table 4.** Two-way ANOVA results for *Wolbachia* transcript abundance. ****p < 0.0001.

| Genus | Term | Df | Sum Sq | Mean<br>Sq | F value | p-value |
| --- | --- | --- | --- | --- | --- | --- |
| *Wolbachia* | Age | 1 | 1351.38 | 1351.38 | 30.71 | <0.0001**<br>** |
|  | Status | 1 | 97.02 | 97.02 | 2.20 | 0.16 |
|  | Age x<br>Status | 1 | 17.23 | 17.23 | 0.39 | 0.54 |
|  | Residual | 13 | 572.14 | 44.01 |  |  |

### 3.5 Age-dependent shifts in transcriptional microbial profiles in *Cx. pipiens*

To further assess age-dependent shift in transcriptional microbiome composition, a two-way ANOVA was performed on the *Cx. pipiens* microbiome, considering only taxa with relative transcript abundance greater than 1%. Our analysis indicated a significant age-related decline in overall transcriptional microbiome composition (p < 0.0001) in both control and infected conditions, suggesting that age may shape the *Culex* transcriptional microbiome (Table 5). However, no significant interaction between infection status and age was observed (p = 0.36). The transcriptional microbiome in this dataset was strongly dominated by *Wolbachia* reads. Therefore, to better evaluate underlying microbial community dynamics, *Wolbachia* sequences were excluded, and an additional analysis was performed. When *Wolbachia* was removed, the analysis still indicated a significant effect of age (p < 0.01) on the remaining microbial taxa (Table 6, Fig. 7). Together, these findings indicate that transcriptional microbiome composition in *Culex* is influenced by host aging primarily through age-dependent increase in *Wolbachia* density, which proportionally reduces relative transcript abundance of other taxa in both control and RVFV-infected mosquitoes (Fig. 7).

**Fig. 7.**
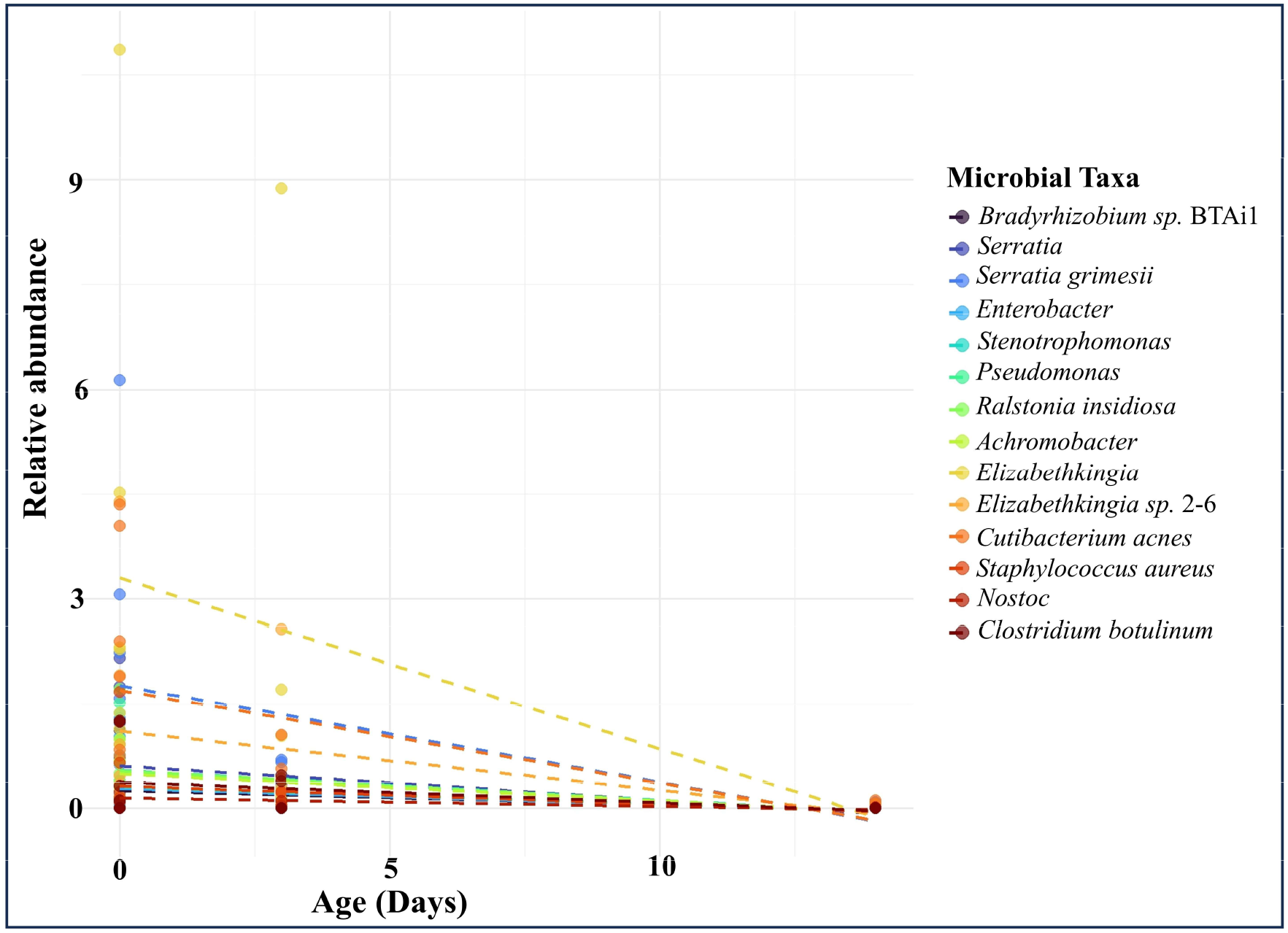
Line plot depicting relative transcript abundance of non-*Wolbachia* microbial taxa across *Culex* age groups (0, 3, and 14 days) over time. This plot focuses on microbial taxa other than *Wolbachia*, with each colored point representing an individual sample’s relative transcript abundance. Dashed linear trend lines illustrate the abundance trajectory of each taxon over time. The overall microbial transcript (excluding *Wolbachia*) appears to diminish in diversity and abundance as mosquitoes mature, suggesting increasing dominance by *Wolbachia* transcripts or suppression of other taxa.

**Table 5.** Two-way ANOVA results for *Culex* transcriptional microbiome data. **p < 0.01, ****p < 0.0001.

| Factor | Df | Sum Sq | Mean Sq | F value | Pr(>F) |
| --- | --- | --- | --- | --- | --- |
| Status | 1 | 0.20 | 0.20 | 9.88 | <0.01** |
| Age | 2 | 1.62 | 0.81 | 39.29 | <0.0001**** |
| Status x Age | 2 | 0.05 | 0.02 | 1.12 | 0.36 |
| Residual | 11 | 0.23 | 0.02 |  |  |

**Table 6.**
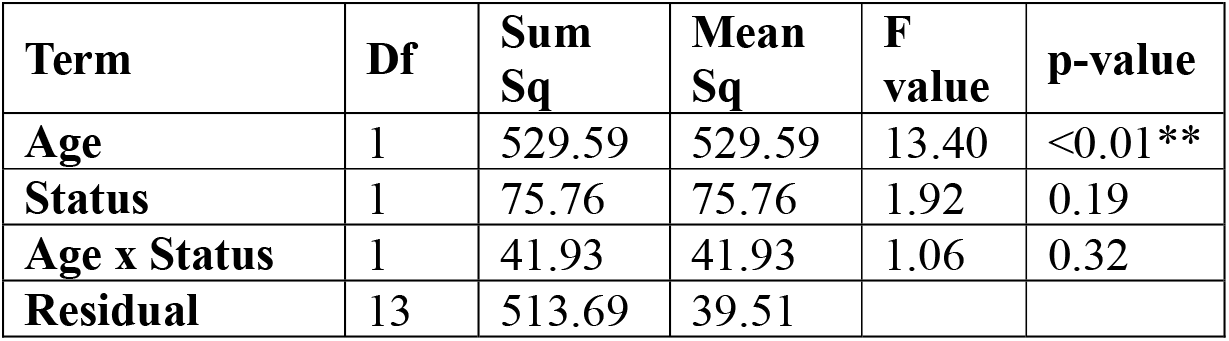
Two-way ANOVA results for Other microbial transcript abundance. **p < 0.01.

| Term | Df | Sum Sq | Mean Sq | F value | p-value |
| --- | --- | --- | --- | --- | --- |
| Age | 1 | 529.59 | 529.59 | 13.40 | <0.01** |
| Status | 1 | 75.76 | 75.76 | 1.92 | 0.19 |
| Age x Status | 1 | 41.93 | 41.93 | 1.06 | 0.32 |
| Residual | 13 | 513.69 | 39.51 |  |  |

To further clarify the infection-associated shifts in microbial transcripts that may be obscured by the high abundance of *Wolbachia* reads, we visualized the composition of non-*Wolbachia* taxa using a log₁₀-scaled heatmap (Supplementary Fig. S1). This representation highlighted that as *Wolbachia* becomes increasingly dominant with age, the relative representation of other bacterial groups decreases, even though their absolute transcript abundances remain low but relatively stable.

## 4. Discussion

The microbiome of a host contributes to essential processes such as nutrient absorption, growth, development, and, most importantly, immune regulation. However, its composition and stability are highly context-dependent, shaped by diet, ecology, and symbiotic partnerships. Pathogenic infections can disrupt this delicate balance, leading to microbiome dysbiosis [52]. Infection-induced microbiome dysbiosis overstimulates the immune system, altering the infection outcome and exacerbating disease pathology [53]. On the other hand, insects can use their microbiome to circumvent the adverse outcomes of pathogenic infections. Several studies have reported microbiome modulation upon bacterial infection, while a few studies have shown that viral infection also leads to changes in the microbiota of insects [54–56].

Our cross-species meta-analysis of publicly accessible datasets, encompassing both systemic and oral infection models, directly examined the frequency and magnitude of viral infection-induced microbiome perturbations at transcriptional level across insect species. After analyzing publicly available host-associated transcriptomics data, we provide preliminary evidence that viral infections are associated with infection-induced shifts in transcriptional microbial community across multiple insect species. Viral infections thus appear to act as powerful drivers of microbiome disruption across insects, but such changes are not universal. Notably, the insects used in our study were reared under different feeding conditions: *Culex* mosquitoes were maintained on a sucrose diet after eclosion and later exposed to infected or uninfected blood meals, aphids were infected via an artificial diet containing crude viral preparation but subsequently maintained on their native host plant (*Vicia villosa*), *A. pernyi* larvae were fed oak foliage, and *Drosophila* were reared on cornmeal agar. Because feeding behavior and gut environment are known to strongly influence microbial community composition, the observed variation in microbiome responses to infection may partly reflect these dietary and ecological differences among hosts. In the absence of complementary laboratory or ecological validation, such host-specific influences remain unresolved; nevertheless, this analysis helps bridge an important gap by providing initial cross-species insights into how symbionts, viruses, and hosts may interact.

Transcriptional microbiome composition of *Wolbachia*-colonized insects (*Cx. pipiens* and *Wolbachia*-positive *D. melanogaster* lines) remained largely unchanged following viral infection, in contrast to marked microbiome perturbations in *Wolbachia*-free host *(D. melanogaster*). This relative stability is consistent with previous studies demonstrating antiviral effects of *Wolbachia* via immune modulation (e.g., ROS, immune pathways), oxidative environment alteration, and reduced viral replication [26,57–59]. While our results do not explicitly show a causative effect of *Wolbachia* on the preservation of the microbial homeostasis, it appears that *Wolbachia* dominance may be related to the increased stability of microbial transcriptome expression profiles in the presence of virus infection. In *Cx. pipiens*, apparent age-associated decline in microbial transcript diversity is primarily driven by rising *Wolbachia* dominance rather than absolute loss of other taxa, consistent with 16S rRNA amplicon studies showing developmental succession toward *Wolbachia* dominance and reduced microbial evenness and diversity in *Culex* [50]. In *A. pisum*, *B. aphidicola* remained the only taxon exceeding 1% relative transcript abundance after infection, consistent with its role as an obligate nutritional symbiont that persists even under viral challenge and reinforcing the idea that essential endosymbionts can maintain stability despite infection-induced stress. More broadly, because our study does not experimentally distinguish obligate intracellular symbionts (e.g., *Wolbachia, Buchnera*) from facultative gut bacteria or transient taxa, apparent microbial transcript differences may reflect a combination of true biological variation and technical read dominance, highlighting the need for caution when interpreting endosymbiont-driven patterns of microbiome stability as we are using RNA-seq datasets.

By extracting unmapped reads from RNA-seq datasets and classifying them with Kraken2, we profiled microbial taxa and characterized transcriptional microbiome composition. Taxonomic classification is limited by the reference database used (Kraken2 PlusPF), and therefore taxa not represented or poorly annotated in the database may not be detected. Variation in sequencing depth and classification rates across samples may influence detection sensitivity; however, the use of relative transcript abundance normalization mitigates these effects and allows comparison across samples. A summary of classified, bacterial, and viral read proportions across datasets is provided in (Supplementary Table S1), highlighting dataset-specific differences in detection sensitivity associated with library preparation strategies. Because our analysis is based on unmapped reads from host RNA-seq datasets, the detected microbial profiles represent transcriptionally active microbial sequences rather than direct measurements of microbial abundance. Therefore, the diversity and relative abundance values reported here should be interpreted as transcript-derived representation of microbial taxa, rather than absolute microbial abundance. This approach revealed transcriptional microbial profile of host that qualitatively align with existing 16S amplicon sequencing studies in these four insects [60–63], thereby supporting the utility of reanalyzing host transcriptomics for microbiota insights [64]. Observed differences in transcriptional microbiome composition among the four insects, as well as some differences in microbiome profile of host between our RNA-seq-derived profiles and 16S datasets, likely reflect host-specific ecological and physiological factors, including diet, feeding habits, gut physiochemistry, developmental stage, and symbiotic associations [3]. Dataset-specific features also have to be taken into account when analyzing this data. The *A. pernyi* dataset was collected using dissected cuticle tissue of larvae, not whole organisms. Thus, the microbial transcript profiles obtained from this dataset reflect tissue-specific responses associated with cuticle liquefaction and may not fully represent microbiome dynamics occurring at the whole-organism level. An additional limitation arises from the use of some poly(A)-selected RNA-seq datasets [33,34]. This approach preferentially captures polyadenylated transcripts, including host mRNA and many viral RNAs, thereby facilitating detection of actively replicating viruses. However, most bacterial transcripts lack poly(A) tails and are therefore likely underrepresented. However, previous studies have shown that this bias does not necessarily lead to a consistent loss of bacterial detection [65]. For example, Melnick et al. (2021) reported comparable recovery of bacterial reads between poly(A)-selected and total RNA-seq datasets, likely due to incomplete stringency of poly(A) selection. Consistent with this, we detected bacterial reads in our analysis, supporting the feasibility of post-hoc microbial profiling from transcriptomic data. Importantly, the RNA-seq datasets analyzed here were generated using different library preparation strategies, which further influence detection sensitivity. In particular, poly(A)-selected datasets may underrepresent or fail to capture viruses that lack polyadenylated transcripts, such as RVFV, whereas viruses with polyadenylated transcripts remain detectable. Additionally, since there were no RVFV-related reads found in the Cp dataset, even though RVFV reference sequences were available in the Kraken2 database, the infection status of these samples could not be validated solely through the RNA-seq data. As a result, findings from this dataset should be interpreted with caution and considered within the framework of the original study, which indicated RVFV exposure and an elevated viral load based on RT-qPCR analyses. As a result, the transcriptional microbial profiles described here may be biased toward polyadenylated organisms and should be interpreted as a partial representation of the microbiome. Nevertheless, the overall taxonomic patterns observed were broadly consistent with previously reported 16S amplicon sequencing studies, supporting the biological relevance of the detected microbial profiles at transcriptome level despite these biases.

Our statistical analyses also underscore the importance of methodological choice when interpreting infection-associated transcriptional microbial changes. PERMANOVA, a commonly used distance-based method, operates on beta-diversity distance matrices and is sensitive to small sample sizes and high within-group variability. In contrast, GLM-based approaches allow testing associations between explanatory variables and diversity metrics such as the Shannon index, complementing distance-based community analyses such as PERMANOVA. Occasional discrepancies between PERMANOVA and GLM results in our datasets are likely due to underlying methodological differences and emphasize that both community composition analyses and diversity-based statistical models are valuable for characterizing infection-associated microbiome shifts.

Our study analyzed existing transcriptomic datasets rather than generating new experimental data, yet it provides complementary insights into how host-microbe-virus dynamics manifest across taxa. By leveraging a meta-transcriptomic perspective, our findings reveal potential links between viral infection and transcriptional microbial variability that may extend beyond *Wolbachia*-associated systems. These patterns suggest that viral-microbial interactions may be more pervasive and functionally significant than currently appreciated. Such patterns offer a foundation for future experimental validation and may help refine our understanding of how immune modulation, symbiont composition, and viral pressures jointly shape host-associated microbial communities. Overall, this study provides a framework for assessing how immune modulation, symbiont composition, host physiology (age), and viral pressures jointly shape host-associated microbiome composition. These findings broaden current perspectives on host-microbe-pathogen interactions and have important implications for insect microbiome research and vector biology, particularly by highlighting the potential to reduce pathogen transmission through management of microbial or symbiont communities.

However, several key questions remain-particularly regarding how *Wolbachia* provides antiviral and microbiome protection, what happens to microbes and viruses during long-term infection or recovery, and how other symbionts or host genes influence these processes. Addressing these gaps will require future studies that integrate controlled infection experiments, ecological metadata, long-term microbiome monitoring, and functional metagenomic analyses to validate and expand these findings.

## 5. Conclusion

The meta-transcriptomic analysis across four insect species indicates that viral infections are commonly associated with transcriptional microbiome perturbations, although the extent and nature of these changes vary substantially among hosts. In *D. melanogaster* and *A. pisum*, viral infection corresponded with a marked reduction in relative microbial transcripts, characterized by decrease in number of detected taxa and dominance of the infecting virus, and a similar pattern was observed in *A. pernyi*, where only the iflavirus persisted post-infection. In contrast, *Cx. pipiens* exhibited remarkable transcriptional microbiome stability following viral infection, with *Wolbachia*-colonized insects showing largely unchanged community composition, suggesting that *Wolbachia* dominance may be associated with transcriptional microbiome stability during viral exposure. Age also emerged as a key determinant of microbial transcript composition in *Cx. pipiens*, with older individuals displaying reduced microbial transcript diversity independent of infection status. This age-dependent microbial transcript succession, together with *Wolbachia* dominance, likely contributes to the observed stability of the microbiome under viral stress in this species. Overall, these findings suggest that transcriptional microbiome stability in insects under viral challenge is shaped by both intrinsic host factors and symbiotic associations, emphasizing the ecological complexity of host-virus-microbiome interactions. This study also demonstrates the potential of re-analyzing host transcriptomic datasets to recover biologically meaningful microbiome signals across diverse insect systems. These insights lay the groundwork for future functional validation of symbiont-mediated viral resilience across insect taxa.

## Author contribution statement

**AM:** conceptualization, data collection and curation, methodology, data analysis and visualization, interpretation and writing–original draft preparation. **TS:** data collection and curation, methodology, data analysis. **VG:** conceptualization, data analysis, interpretation, writing–reviewing and editing, supervision and funding acquisition. All authors read and approved the final manuscript.

## Data Availability Statement

All datasets used in this study are publicly available in the NCBI SRA under BioProject IDs PRJNA682591, PRJNA596664, PRJNA419454, and PRJNA517653. Additional analysis scripts used in this study will be made publicly available in a dedicated repository (https://github.com/AdwaitMishra/Metatranscriptomic-Analysis).

## Acknowledgements

Not applicable

## Ethics approval and consent to participate

Not applicable

## Consent for publication

Not applicable

## References

1. Gupta A, Nair S. Dynamics of Insect-Microbiome Interaction Influence Host and Microbial Symbiont. Front Microbiol. 2020;11:1357. 10.3389/fmicb.2020.01357

2. Fredensborg BL, Fossdal Í Kálvalíð I, Johannesen TB, Stensvold CR, Nielsen HV, Kapel CMO. Parasites modulate the gut-microbiome in insects: A proof-of-concept study. PLoS One. Public Library of Science (PLoS); 2020;15:e0227561. 10.1371/journal.pone.0227561

3. Engel P, Moran NA. The gut microbiota of insects - diversity in structure and function. FEMS Microbiol Rev. Oxford University Press (OUP); 2013;37:699–735. 10.1111/1574-6976.12025

4. Ramalho MO, Bueno OC, Moreau CS. Species-specific signatures of the microbiome from Camponotus and Colobopsis ants across developmental stages. PLoS One. 2017;12:e0187461. 10.1371/journal.pone.0187461

5. Yuan J, Chaparro JM, Manter DK, Zhang R, Vivanco JM, Shen Q. Roots from distinct plant developmental stages are capable of rapidly selecting their own microbiome without the influence of environmental and soil edaphic factors. Soil Biol Biochem. Elsevier BV; 2015;89:206–9. 10.1016/j.soilbio.2015.07.009

6. Salazar N, Valdés-Varela L, González S, Gueimonde M, de Los Reyes-Gavilán CG. Nutrition and the gut microbiome in the elderly. Gut Microbes. 2017;8:82–97. 10.1080/19490976.2016.1256525

7. Yatsunenko T, Rey FE, Manary MJ, Trehan I, Dominguez-Bello MG, Contreras M, et al. Human gut microbiome viewed across age and geography. Nature. Springer Science and Business Media LLC; 2012;486:222–7. 10.1038/nature11053

8. Hacquard S, Spaepen S, Garrido-Oter R, Schulze-Lefert P. Interplay between innate immunity and the plant Microbiota. Annu Rev Phytopathol. Annual Reviews; 2017;55:565–89. 10.1146/annurev-phyto-080516-035623

9. Taylor M, Vega NM. Host immunity alters community ecology and stability of the microbiome in a Caenorhabditis elegans model. mSystems. American Society for Microbiology; 2021;6. 10.1128/mSystems.00608-20

10. Marra A, Hanson MA, Kondo S, Erkosar B, Lemaitre B. Drosophila antimicrobial peptides and lysozymes regulate gut Microbiota composition and abundance. MBio. American Society for Microbiology; 2021;12:e0082421. 10.1128/mBio.00824-21

11. Hanada S, Pirzadeh M, Carver KY, Deng JC. Respiratory viral infection-induced microbiome alterations and secondary bacterial pneumonia. Front Immunol. Frontiers Media SA; 2018;9:2640. 10.3389/fimmu.2018.02640

12. Groves HT, Higham SL, Moffatt MF, Cox MJ, Tregoning JS. Respiratory viral infection alters the gut Microbiota by inducing inappetence. MBio. American Society for Microbiology; 2020;11. 10.1128/mbio.03236-19

13. Zhang F, Weckhorst JL, Assié A, Hosea C, Ayoub CA, Khodakova AS, et al. Natural genetic variation drives microbiome selection in the Caenorhabditis elegans gut. Curr Biol. Elsevier BV; 2021;31:2603–2618.e9. 10.1016/j.cub.2021.04.046

14. Champion CJ, Xu J. The impact of metagenomic interplay on the mosquito redox homeostasis. Free Radic Biol Med. 2017;105:79–85. 10.1016/j.freeradbiomed.2016.11.031

15. Broderick NA, Lemaitre B. Gut-associated microbes of Drosophila melanogaster. Gut Microbes. 2012;3:307–21. 10.4161/gmic.19896

16. Wong CNA, Ng P, Douglas AE. Low-diversity bacterial community in the gut of the fruitfly Drosophila melanogaster. Environ Microbiol. Wiley; 2011;13:1889–900. 10.1111/j.1462-2920.2011.02511.x

17. Feng H, Edwards N, Anderson CMH, Althaus M, Duncan RP, Hsu Y-C, et al. Trading amino acids at the aphid-Buchnera symbiotic interface. Proc Natl Acad Sci U S A. Proceedings of the National Academy of Sciences; 2019;116:16003–11. 10.1073/pnas.1906223116

18. Wang H, Zhang J-Y, Wang X-M, Hu H-L, Xia R-X, Li Q, et al. Comparison of bacterial communities between midgut and midgut contents in two silkworms, Antheraea pernyi and Bombyx mori. Sci Rep. Springer Science and Business Media LLC; 2020 [cited 2025 Dec 17];10:12966. 10.1038/s41598-020-69906-y

19. Hou P, Liu L, Ma X, Men Y, Yang D, Wang J, et al. Diversity characteristics and composition of gut Microbiota in Antheraea pernyi (Lepidoptera: Saturniidae) larvae across different instars. Insects. MDPI AG; 2025;16:909. 10.3390/insects16090909

20. Douglas AE. Multiorganismal insects: diversity and function of resident microorganisms. Annu Rev Entomol. Annual Reviews; 2015;60:17–34. 10.1146/annurev-ento-010814-020822

21. Vodovar N, Vinals M, Liehl P, Basset A, Degrouard J, Spellman P, et al. Drosophila host defense after oral infection by an entomopathogenic Pseudomonas species. Proc Natl Acad Sci U S A. Proceedings of the National Academy of Sciences; 2005;102:11414–9. 10.1073/pnas.0502240102

22. Marasco R, Fusi M, Callegari M, Jucker C, Mapelli F, Borin S, et al. Destabilization of the Bacterial Interactome Identifies Nutrient Restriction-Induced Dysbiosis in Insect Guts. Microbiol Spectr. 2022;10:e0158021. 10.1128/spectrum.01580-21

23. Johnson KN. Bacteria and antiviral immunity in insects. Curr Opin Insect Sci. Elsevier BV; 2015;8:97–103. 10.1016/j.cois.2015.01.008

24. Cansado-Utrilla C, Zhao SY, McCall PJ, Coon KL, Hughes GL. The microbiome and mosquito vectorial capacity: rich potential for discovery and translation. Microbiome. 2021;9:111. 10.1186/s40168-021-01073-2

25. Wei G, Lai Y, Wang G, Chen H, Li F, Wang S. Insect pathogenic fungus interacts with the gut microbiota to accelerate mosquito mortality. Proc Natl Acad Sci U S A. 2017;114:5994–9. 10.1073/pnas.1703546114

26. Simhadri RK, Fast EM, Guo R, Schultz MJ, Vaisman N, Ortiz L, et al. The gut commensal microbiome of Drosophila melanogaster is modified by the Endosymbiont Wolbachia. mSphere. American Society for Microbiology; 2017;2. 10.1128/mSphere.00287-17

27. Ye YH, Seleznev A, Flores HA, Woolfit M, McGraw EA. Gut microbiota in Drosophila melanogaster interacts with Wolbachia but does not contribute to Wolbachia -mediated antiviral protection. J Invertebr Pathol. Elsevier BV; 2017;143:18–25. 10.1016/j.jip.2016.11.011

28. Pfeiffer JK, Virgin HW. Viral immunity. Transkingdom control of viral infection and immunity in the mammalian intestine. Science. American Association for the Advancement of Science (AAAS); 2016;351:aad5872–aad5872. 10.1126/science.aad5872

29. Duerkop BA, Hooper LV. Resident viruses and their interactions with the immune system. Nat Immunol. Springer Science and Business Media LLC; 2013 [cited 2025 May 29];14:654–9. 10.1038/ni.2614

30. Kernbauer E, Ding Y, Cadwell K. An enteric virus can replace the beneficial function of commensal bacteria. Nature. Springer Science and Business Media LLC; 2014 [cited 2025 May 29];516:94–8. 10.1038/nature13960

31. Robinson CM, Pfeiffer JK. Viruses and the Microbiota. Annu Rev Virol. Annual Reviews; 2014;1:55–69. 10.1146/annurev-virology-031413-085550

32. Lindsey ARI, Bhattacharya T, Hardy RW, Newton ILG. *Wolbachia* and virus alter the host transcriptome at the interface of nucleotide metabolism pathways. MBio. American Society for Microbiology; 2021;12. 10.1128/mbio.03472-20

33. Núñez AI, Esteve-Codina A, Gómez-Garrido J, Brustolin M, Talavera S, Berdugo M, et al. Alteration in the Culex pipiens transcriptome reveals diverse mechanisms of the mosquito immune system implicated upon Rift Valley fever phlebovirus exposure. PLoS Negl Trop Dis. 2020;14:e0008870. 10.1371/journal.pntd.0008870

34. Wang G, Na S, Duan X, Leng Z, Jiang Y, Shi S, et al. Transcriptome sequencing to unravel the molecular mechanisms underlying the cuticle liquefaction of Antheraea pernyi following Antheraea pernyi nucleopolyhedrovirus challenge. Mol Immunol. 2019;109:108–15. 10.1016/j.molimm.2019.03.009

35. Lu H, Zhu J, Yu J, Chen X, Kang L, Cui F. A symbiotic virus facilitates aphid adaptation to host plants by suppressing jasmonic acid responses. Mol Plant Microbe Interact. Scientific Societies; 2020;33:55–65. 10.1094/MPMI-01-19-0016-R

36. Abbà S, Rossi M, Vallino M, Galetto L, Marzachì C, Turina M. Metatranscriptomic Assessment of the Microbial Community Associated With the Flavescence dorée Phytoplasma Insect Vector *Scaphoideus titanus*. Front Microbiol. Frontiers Media SA; 2022;13:866523. 10.3389/fmicb.2022.866523

37. Espín-Sánchez D, Russo L, Diambra L, Mannino MC, Scorsetti AC, Huarte-Bonnet C, et al. Host transcriptional and microbiome metatranscriptomic changes in soybean plants carrying the insect-pathogenic fungus Beauveria bassiana as an endophyte. J Invertebr Pathol. Elsevier BV; 2026;217:108629. 10.1016/j.jip.2026.108629

38. Xu Q, Zhang Y, Tao S. Functional dynamics reveal the response of the crabapple (Malus sp.) phyllosphere microbiome to Gymnosporangium yamadae infection. mSystems. American Society for Microbiology; 2025;10:e0084325. 10.1128/msystems.00843-25

39. Andrews S, Others. FastQC: a quality control tool for high throughput sequence data. Babraham Bioinformatics, Babraham Institute, Cambridge, United Kingdom; 2010.

40. Chen S, Zhou Y, Chen Y, Gu J. fastp: an ultra-fast all-in-one FASTQ preprocessor. Bioinformatics. 2018;34:i884–90. 10.1093/bioinformatics/bty560

41. Dobin A, Davis CA, Schlesinger F, Drenkow J, Zaleski C, Jha S, et al. STAR: ultrafast universal RNA-seq aligner. Bioinformatics. 2013;29:15–21. 10.1093/bioinformatics/bts635

42. Rajagopala SV, Bakhoum NG, Pakala SB, Shilts MH, Rosas-Salazar C, Mai A, et al. Metatranscriptomics to characterize respiratory virome, microbiome, and host response directly from clinical samples. Cell Rep Methods. Elsevier BV; 2021;1:100091. 10.1016/j.crmeth.2021.100091

43. Li H, Handsaker B, Wysoker A, Fennell T, Ruan J, Homer N, et al. The Sequence Alignment/Map format and SAMtools. Bioinformatics. Oxford University Press (OUP); 2009;25:2078–9. 10.1093/bioinformatics/btp352

44. Bushnell B, Rood J, Singer E. BBMerge - Accurate paired shotgun read merging via overlap. PLoS One. 2017;12:e0185056. 10.1371/journal.pone.0185056

45. Wood DE, Salzberg SL. Kraken: ultrafast metagenomic sequence classification using exact alignments. Genome Biol. 2014;15:R46. 10.1186/gb-2014-15-3-r46

46. Johnson KV-A, Burnet PWJ. Microbiome: Should we diversify from diversity? Gut Microbes. 2016;7:455–8. 10.1080/19490976.2016.1241933

47. Oksanen J, Blanchet FG, Kindt R, Legendre P. vegan: Community Ecology Package. R package version 2.0-10. 2013. 2015; https://scholar.google.com/citations?user=2WBRFVIAAAAJ&hl=en&oi=sra

48. Wickham H. Ggplot2. Wiley Interdiscip Rev Comput Stat. Wiley; 2011;3:180–5. 10.1002/wics.147

49. Kolde R, Kolde MR. Package ‘pheatmap. 2015;1.

50. Flores GAM, Lopez RP, Cerrudo CS, Perotti MA, Consolo VF, Berón CM. Wolbachia dominance influences the Culex quinquefasciatus microbiota. Sci Rep. Springer Science and Business Media LLC; 2023;13:18980. 10.1038/s41598-023-46067-2

51. Duguma D, Hall MW, Rugman-Jones P, Stouthamer R, Terenius O, Neufeld JD, et al. Developmental succession of the microbiome of Culex mosquitoes. BMC Microbiol. Springer Science and Business Media LLC; 2015;15. 10.1186/s12866-015-0475-8

52. Pham TAN, Lawley TD. Emerging insights on intestinal dysbiosis during bacterial infections. Curr Opin Microbiol. Elsevier BV; 2014;17:67–74. 10.1016/j.mib.2013.12.002

53. Clemente JC, Ursell LK, Parfrey LW, Knight R. The impact of the gut microbiota on human health: an integrative view. Cell. Elsevier BV; 2012;148:1258–70. 10.1016/j.cell.2012.01.035

54. Martínez-Solís M, Collado MC, Herrero S. Influence of Diet, Sex, and Viral Infections on the Gut Microbiota Composition of Spodoptera exigua Caterpillars. Front Microbiol. 2020;11:753. 10.3389/fmicb.2020.00753

55. Zouache K, Michelland RJ, Failloux A-B, Grundmann GL, Mavingui P. Chikungunya virus impacts the diversity of symbiotic bacteria in mosquito vector. Mol Ecol. 2012;21:2297–309. 10.1111/j.1365-294X.2012.05526.x

56. Yongsawas R, Chaimanee V, Pettis JS, Boncristiani Junior HF, Lopez D, In-On A, et al. Impact of Sacbrood Virus on Larval Microbiome of Apis mellifera and Apis cerana. Insects. 2020;11. 10.3390/insects11070439

57. Bruner-Montero G, Jiggins FM. Wolbachia protects Drosophila melanogaster against two naturally occurring and virulent viral pathogens. Sci Rep. 2023;13:8518. 10.1038/s41598-023-35726-z

58. Cogni R, Ding SD, Pimentel AC, Day JP, Jiggins FM. Wolbachia reduces virus infection in a natural population of Drosophila. Commun Biol. Springer Science and Business Media LLC; 2021 [cited 2023 Sept 19];4:1–7. 10.1038/s42003-021-02838-z

59. Zug R, Hammerstein P. Wolbachia and the insect immune system: what reactive oxygen species can tell us about the mechanisms of Wolbachia-host interactions. Front Microbiol. Frontiers Media SA; 2015;6:1201. 10.3389/fmicb.2015.01201

60. Cho KH, Kang SO. The gut Microbiota of Drosophila melanogaster: A model for host-microbe interactions in metabolism, immunity, behavior, and disease. Microorganisms. MDPI AG; 2025;13:2515. 10.3390/microorganisms13112515

61. Allahem A, Alajmi R, Alajami MA, Al-Ghamdi S, Bashir MA, Zeb J, et al. Molecular Detection of natural bacterial flora within Culex pipiens pipiens (Diptera: Culicidae) in Aleppo, Syria. J Genet Eng Biotechnol. Elsevier BV; 2025;23:100604. 10.1016/j.jgeb.2025.100604

62. Gauthier J-P, Outreman Y, Mieuzet L, Simon J-C. Bacterial communities associated with host-adapted populations of pea aphids revealed by deep sequencing of 16S ribosomal DNA. PLoS One. 2015;10:e0120664. 10.1371/journal.pone.0120664

63. Xu X, Gao Y, Ren S, Liu Z, Zhang Y, Zhang Z, et al. An analysis of the gut Microbiota of fifth-instar Antheraea pernyi larvae and a functional exploration of a Bacillus subtilis strain. Insects. MDPI AG; 2025;16:333. 10.3390/insects16040333

64. Leimena MM, Ramiro-Garcia J, Davids M, van den Bogert B, Smidt H, Smid EJ, et al. A comprehensive metatranscriptome analysis pipeline and its validation using human small intestine microbiota datasets. BMC Genomics. Springer Science and Business Media LLC; 2013;14:530. 10.1186/1471-2164-14-530

65. Melnick M, Gonzales P, LaRocca TJ, Song Y, Wuu J, Benatar M, et al. Application of a bioinformatic pipeline to RNA-seq data identifies novel virus-like sequence in human blood. G3 (Bethesda). Oxford University Press (OUP); 2021;11. 10.1093/g3journal/jkab141

