## Supplementary material for "A cross-species meta-transcriptomic analysis of viral infection-induced shifts in microbial transcript profiles across insects"

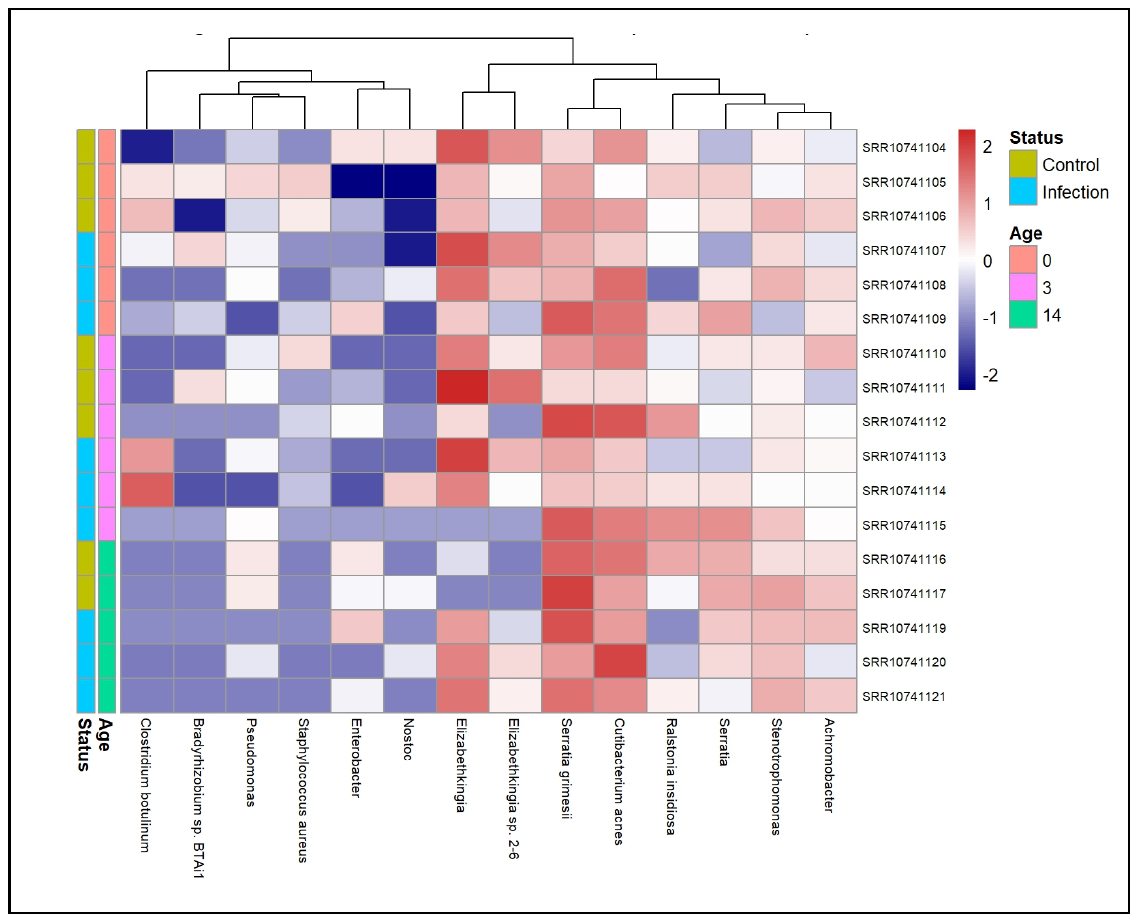
**Supplementary material**

**Fig. S1** Heatmap of log₁₀-transformed raw transcript counts of non-*Wolbachia* bacterial taxa selected based on >1% relative transcript abundance across Cp samples. Columns represent samples annotated by age and infection status, and rows represent taxa. Values were log₁₀-transformed (log₁₀[count + 1]) and scaled by row (z-score) to highlight relative differences across samples.

**Table S1 Proportion of classified, bacterial, and viral reads across RNA-seq datasets. Values represent the mean percentage of reads assigned to microbial taxa (classified reads), as well as bacterial and viral fractions, in control and infected samples for each dataset, based on Kraken2 classification of non-host reads.** Percentages were calculated relative to total non-host reads per sample and averaged across replicates within each condition. The table also indicates whether datasets were generated using poly(A)-selected or non-poly(A) library preparation. Variation in read proportions reflects both biological differences and technical factors, including library preparation strategy and sequencing depth, which may influence detection sensitivity. Notably, classified reads include assignments across multiple taxonomic domains; therefore, higher classification percentages do not necessarily correspond exclusively to bacterial or viral reads.

|  | **Poly-A** | **Classified** | | **Bacteria** | | **Virus** | |
| --- | --- | --- | --- | --- | --- | --- | --- |
|  |  | Control | Infected | Control | Infected | Control | Infected |
| ***D. melanogaster*** | No | 6.69 | 11.94 | 1.79 | 0.60 | 1.22 | 7.14 |
| ***Cx. pipiens*** | Yes | 4.95 | 7.43 | 0.52 | 0.79 | 0.00 | 0.00 |
| ***A. pisum*** | No | 58.52 | 42.25 | 4.59 | 0.61 | 0.55 | 34.53 |
| ***A. pernyi*** | Yes | 2.81 | 96.44 | 1.82 | 0.01 | 0.26 | 96.38 |
